# Distinct chromatin signatures of DNA hypomethylation in aging and cancer

**DOI:** 10.1101/229476

**Authors:** Raúl F. Pérez, Juan Ramón Tejedor, Gustavo F. Bayón, Agustín F. Fernández, Mario F. Fraga

## Abstract

**Background:** Cancer is an aging-associated disease but the underlying molecular links between these processes are still largely unknown. Gene promoters that become hypermethylated in aging and cancer share a common chromatin signature in ES cells. In addition, there is also global DNA hypomethylation in both processes. However, any similarities of the regions where this loss of DNA methylation occurs is currently not well characterized, nor is it known whether such regions also share a common chromatin signature in aging and cancer.

**Results:** To address this issue we analysed TCGA DNA methylation data from a total of 2,311 samples, including control and cancer cases from patients with breast, kidney, thyroid, skin, brain and lung tumors and healthy blood, and integrated the results with histone, chromatin state and transcription factor binding site data from the NIH Roadmap Epigenomics and ENCODE projects. We identified 98,857 CpG sites differentially methylated in aging, and 286,746 in cancer. Hyper- and hypomethylated changes in both processes each had a similar genomic distribution across tissues and displayed tissue-independent alterations. The identified hypermethylated regions in aging and cancer shared a similar bivalent chromatin signature. In contrast, hypomethylated DNA sequences occurred in very different chromatin contexts. DNA hypomethylated sequences were enriched at genomic regions marked with the activating histone posttranslational modification H3K4me1 in aging, whilst in cancer, loss of DNA methylation was primarily associated with the repressive H3K9me3 mark.

**Conclusions:** Our results suggest that the role of DNA methylation as a molecular link between aging and cancer is more complex than previously thought.

## Background

Age is amongst the most important risk factors for cancer [1-3]. However, the underlying molecular mechanisms governing this relationship are still poorly understood. Recent research has established polycomb-target gene promoter hypermethylation as a common epigenetic characteristic of cancer [4,5]. In this scenario, prior to alteration these promoters display an embryonic stem cell “bivalent chromatin pattern” consisting of the repressive histone mark H3K27me3 and the active mark H3K4me3 [6]. Genes affected by this process are associated with developmental regulation [7], implying a possible stem cell origin of cancer whereby aberrant hypermethylation could promote a continuously self-renewing embryonic-like state in cancer cells. Furthermore, complementary studies have revealed that these bivalent domains still remain following drug induced gene reactivation and that they can spread over adjacent CpG islands [8].

Interestingly, promoter hypermethylation of polycomb-target genes was later described in aging blood [9,10] and other tissue types such as mesenchymal stem cells [11], ovary [10], brain [12,13], kidney and skeletal muscle [13]. These findings, which were also confirmed using whole genome bisulfite sequencing [14], suggest that aging may predispose to cancer by irreversibly perpetuating a stem cell-like status [10].

In addition to aberrant locus-specific DNA hypermethylation, tumoral cells are also globally hypomethylated as compared to their healthy counterparts [15]. While this molecular alteration preferentially occurs at gene bodies, intergenic DNA regions and repeated DNA elements [16,17] and is proposed to be associated with chromosomal instability, reactivation of transposable elements and loss of genomic imprinting, its precise functional role in cancer development is still poorly understood [18]. Intriguingly, global loss of genomic DNA methylation has also been reported during the aging process [19–21]. Whole-genome bisulfite sequencing and methylation arrays have confirmed the global loss of DNA methylation in different human tissues including blood [14], mesenchymal stem cells and brain [11]. On the other hand, other important tissues such as skeletal muscle do not seem to become hypomethylated with aging [22].

Despite the interesting parallelism in aging and cancer recently reported with respect to hypermethylated DNA regions, the relationship between hypomethylated DNA sequences in these two processes has not been systematically studied. To address this issue, here we have analysed DNA methylation changes and their associated chromatin patterns in a total of more than 2,300 healthy and tumoral samples obtained from differentially aged individuals, using HumanMethylation450 BeadChip data generated by The Cancer Genome Atlas (TCGA) consortium and other datasets [23–25]. Our results confirmed the relationship between DNA hypermethylation in aging and in cancer, but they also revealed important differences in DNA hypomethylation changes in the two processes that might be important in order to understand the possible role of DNA methylation as a molecular link between decline related to aging and tumor development.

## Results

### DNA methylation profiling in aging and cancer

To identify DNA methylation changes in aging and cancer we collected DNA methylation data obtained with the HumanMethylation450 BeadChip (Illumina) (see Methods) and compared the DNA methylation status of a total of 361,698 CpG sites across 1,762 samples. These samples corresponded to healthy and tumoral tissues obtained from differentially aged patients with breast, kidney, thyroid, skin and brain tumors (see Table 1 and Additional file 2: Table S1 for extended information).

**Table 1.**
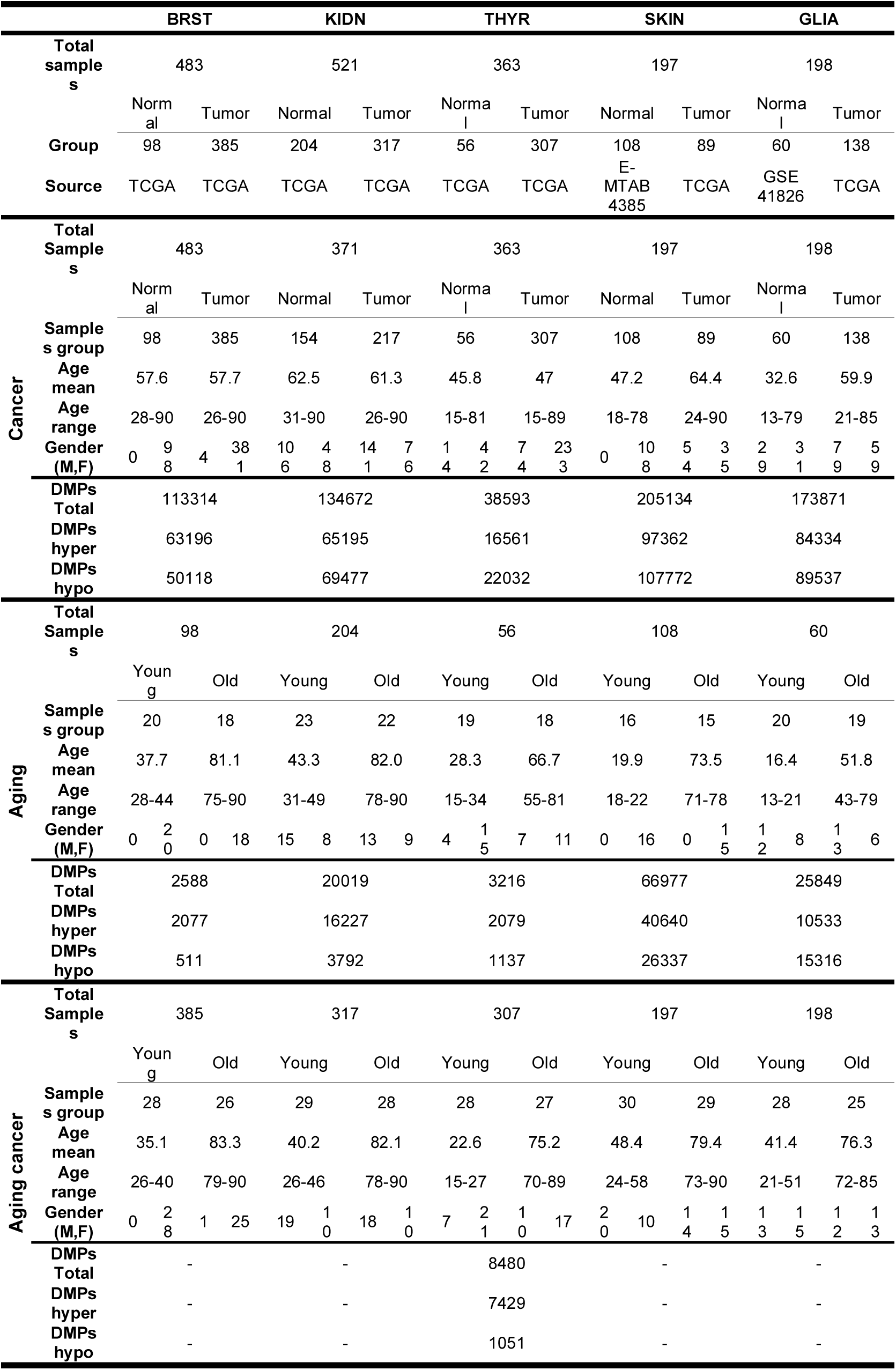
*Description of sample groups and dmCpGs obtained in the analyses.*

Using an empirical Bayes moderated t-test (see Methods), we identified 113,314 autosomal CpG sites that were differentially methylated (dmCpGs; FDR < 0.05) between breast tumor samples and their corresponding non-tumorigenic counterparts (Fig. 1a). Using a similar approach, we identified 134,672 dmCpGs in tumors from kidney, 38,593 from thyroid, 205,143 from skin and 173,871 from brain tumors (Table 1; Fig. 1a, top panel and Additional file 3: Table S2). Hierarchical clustering of samples using the tumor-type specific dmCpGs enabled us to distinguish between tumoral and control samples (Fig. 1b, upper panel; Additional file 1: Figure S1). Although the number of significant hyper- and hypomethylated probes was somewhat variable across the different tumor datasets, the magnitude of change was generally similar and tumor-type independent, with no dominance of either hyper- or hypomethylation changes.

**Fig. 1.**
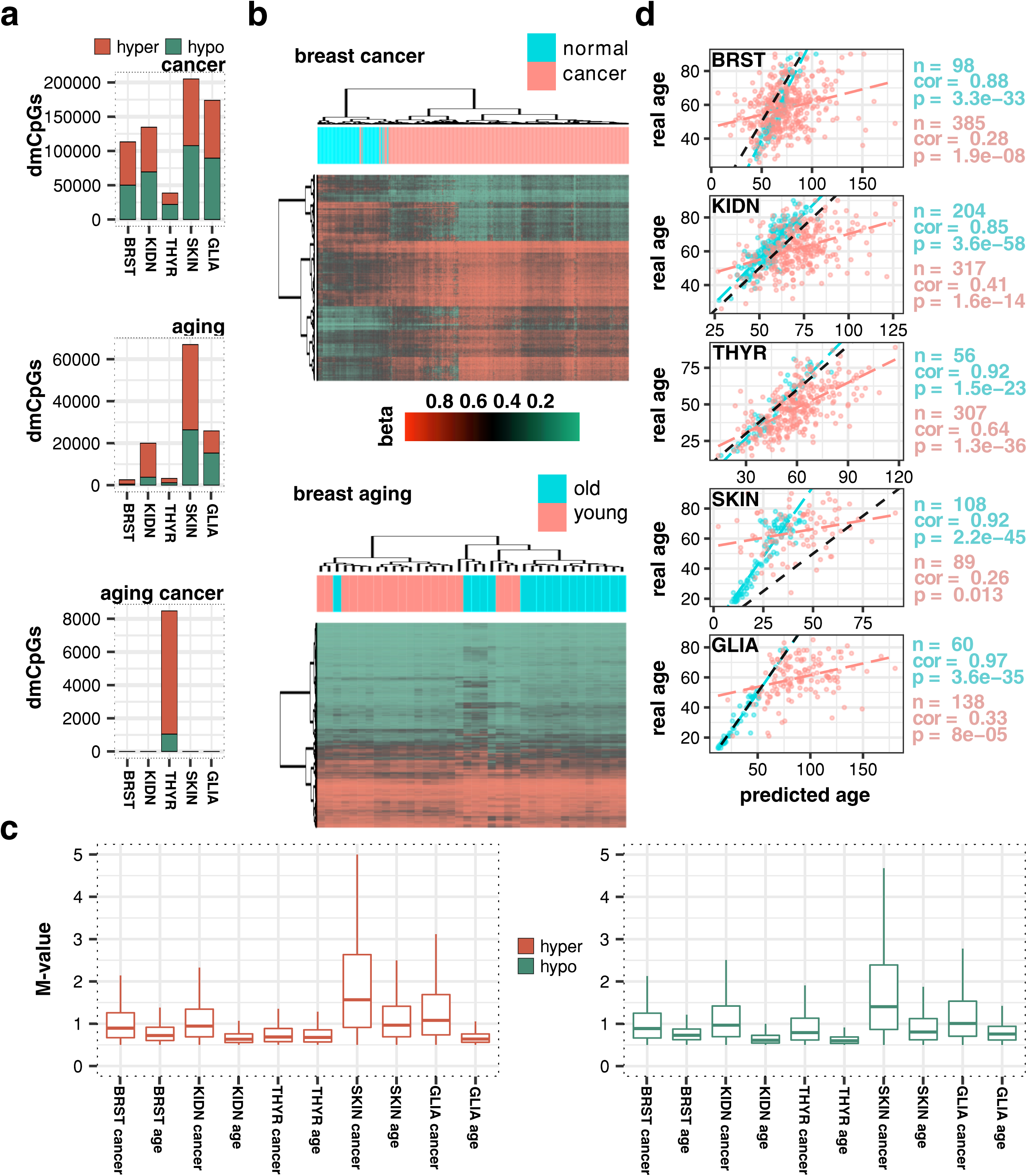
*DNA methylation changes in aging and cancer. **a** Stacked barplots indicating total number of dmCpGs detected in cancer, aging and aging cancer tissues. **b** Hierarchical clustering and heatmaps including the 1,000 most significant dmCpGs for breast cancer and aging analyses. Beta-values of DNA methylation are displayed from zero (green) to one (red). **c** Boxplots comparing magnitude of M-values of methylation changes in cancer and aging. All differences are statistically significant (Wilcoxon tests, all P<0.05, Additional file 4: Table S3). **d** Scatterplots indicating correlation of chronological age with Horvath’s predicted age in normal and cancer samples. Pearson product-moment correlation coefficient (cor) is indicated and linear fit lines are added to help with data interpretation.*

We next studied age-associated DNA methylation changes in the same set of tissue samples. In control cases, an empirical Bayes moderated t-test identified 2,588 autosomal dmCpG sites (FDR < 0.05) between non-tumorigenic breast tissue obtained from young (ages ranging from 28-44 yrs) and elderly (aged between 75 and 90 yrs) donors. Using a similar approach, we found 20,019 dmCpG sites in kidney tissues, 3,216 in equivalent thyroid tissues, 66,977 in skin and 25,849 in brain tissues, obtained from young and elderly individuals (see Table 1 for age ranges; Fig. 1a, middle panel and Additional file 3: Table S2 for additional information). Hierarchical clustering of samples using the tissue-type specific dmCpGs enabled us to classify most of the tissues according to their corresponding age groups (Fig. 1b, lower panel; Additional file 1: Figure S1), but the efficiency of classification was reduced as compared to the clustering obtained with the cancer dmCpGs.

On the whole, and in contrast to cancer, the number of age-associated DNA methylation changes was highly variable and also tissue-type dependent, and the dominant direction of the changes observed tended towards DNA hypermethylation.

Globally, methylation changes were found to be more pronounced in cancer than in aging (Fig. 1c and Additional file 4: Table S3, Wilcoxon tests; all *P*<0.05), while comparison of hyper-versus hypomethylation changes was variable and disease and tissue-type dependent. Intriguingly, with the exception of thyroid cancer, most tumors obtained from differentially-aged patients did not show significant age-associated DNA methylation changes (Table 1, Fig. 1a, bottom panel and Additional file 3: Table S2). Further analyses of normal and cancer samples with Horvath’s predictor (implementation in wateRmelon R package) revealed that thyroid cancer had the highest correlation of real-versus-predicted age across all the cancer types in our dataset (Fig. 1d). This result, which is in the same vein as previous observations [26], suggests that thyroid tumors might be associated with a more conventional age-related progression.

### Genomic distribution of dmCpGs in aging and cancer

The study of the genomic distribution of the dmCpGs revealed that hypomethylated CpG sites followed a similar disease and tissue-independent trend, being preferentially found at low density CpG DNA regions interrogated by the array in both cancer and aging processes (average median difference compared to array 49%, Wilcoxon tests all *P*<0.001) (Fig. 2a; see also Additional file 5: Table S4). Consequently, with respect to the array, these hypomethylated CpG sites were enriched at open sea locations (Fisher’s tests; all *P*<0.001, all odds ratios (ORs) >2.25) (Fig. 2b; Additional file 1: Figure S2 and Additional file 6: Table S5), and intronic and intergenic regions (Fisher’s tests; all *P*<0.001, all ORs>1.34 and >1.21, respectively, except non-significant thyroid aging) (Fig. 2c; Additional file 1: Figure S2 and Additional file 6: Table S5) while impoverished at CpG islands (Fisher’s tests; all *P*<0.001, all ORs<0.43) and gene promoters (Fisher’s tests; all *P*<0.001, all ORs<0.64). Density of hypermethylated CpG sites in cancer was variable but comparable to background array density (average median difference <±12 %, Wilcoxon tests; all *P*<0.001) (Fig. 2a; Additional file 5: Table S4), whereas a noticeably high CpG density was found for breast, kidney and thyroid in the aging context (average median difference 44%, Wilcoxon tests; all *P*<0.001). Consequently, these dmCpGs were enriched at CpG islands (Fisher’s tests; all *P*<0.001, all ORs>1.89) and gene promoters (Fisher’s tests; all *p*<0.001, all ORs>1.08) (Fig. 2c; Additional file 1: Figure S2 and Additional file 6: Table S5).

**Fig. 2.**
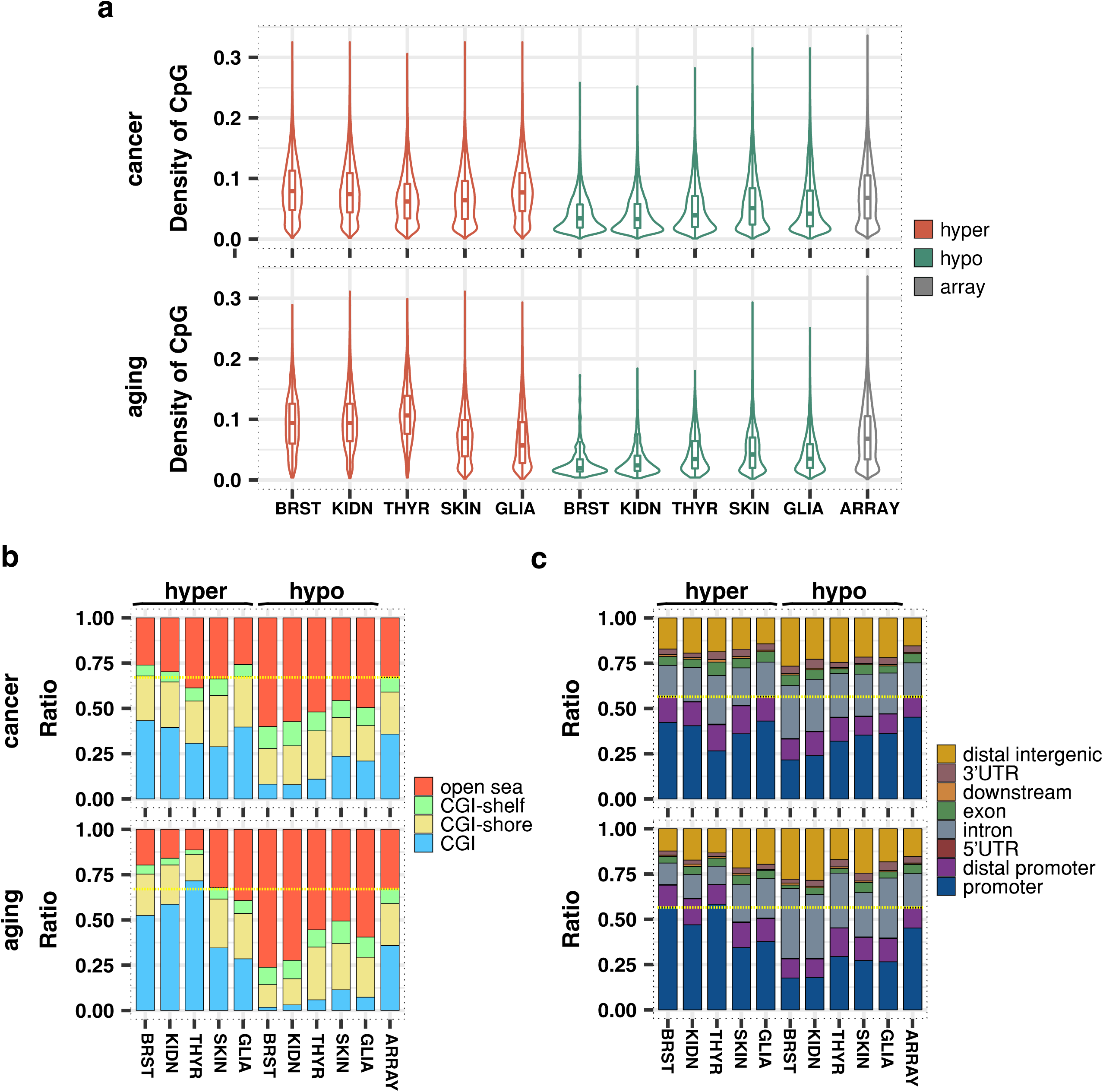
*Similarities and disparities in genomic distribution of methylation changes in aging and cancer. **a** Violin plots showing the distribution of CpG density for cancer and aging hyper- and hypomethylated dmCpGs and the background sites in the Infinium HumanMethylation450 microarray. **b** Stacked barplots indicating relative distribution of differentially methylated CpGs according to their CpG island status. A dashed yellow line separates CpG island associated locations from open sea. **c** Stacked barplots indicating relative distribution of differentially methylated CpGs according to their gene location status. A dashed yellow line separates promoter associated locations from the rest.*

Comparative enrichment analysis confirmed that DNA hypomethylation in aging and cancer mainly occurred at low CpG density DNA regions located at introns, open sea and intergenic DNA regions, while hypermethylation distribution was more irregular and more similar to array distribution. Nonetheless, a common and strong tendency was found when comparing hyper- to hypomethylation changes in both aging and cancer, whereby hypermethylation changes always occurred in regions with a higher CpG density than did hypomethylation changes (average median difference 51% Wilcoxon tests; all *P*<0.001)(Fig. 2a; Additional file 5: Table S4). Accordingly, this resulted in strong differences in local enrichments at CpG islands and open sea locations, as well as gene promoters and intergenic regions, in most cases (Fig. 2b and Fig. 2c; Additional file 1: Figure S3 and Additional file 6: Table S5), with this effect being even more pronounced for the changes accumulated over the aging process.

### Tissue type-independent DNA methylation changes in aging and cancer

To determine the effect of tissue-type on the DNA methylation changes during aging and cancer we compared the previously identified dmCpG sites in breast, kidney, thyroid, skin and brain samples. In cancer, 89,879 (49%) of hyper- and 92,031 (49%) of hypomethylated CpGs were common to at least two different tumor-types (Fig. 3a). Furthermore, 1,962 hypermethylated (1.1%) and 2,708 hypomethylated CpG sites (1.5%) were common to all five tumor-types analysed (Fig. 3b; Additional file 7: Table S6). In contrast, the extent of the overlap between dmCpGs in aging across different tissues was considerably reduced. Indeed, only 10,425 (18%) of the hyper- and 3,366 (8%) of the hypomethylated CpG sites were common to at least two tissue-types and only 89 (0.15%) hyper- and 1 (0.002%) hypomethylated CpG sites were common to all five tissue-types analysed (Fig. 3a and 3b; Additional file 7: Table S6). However, statistical analyses of the pairwise overlaps between the different sets of probes showed overall enrichments in every case, especially for aging (Fig 3d, Fisher’s tests; all *P*<0.001, Additional file 8: Table S7). This over-enrichment was also revealed through a simulation of random sampling of probes from the array (Additional file 1: Figure S4a). Taken together, these results suggest that both cancer and aging manifest tissue independent changes in DNA methylation.

**Fig. 3.**
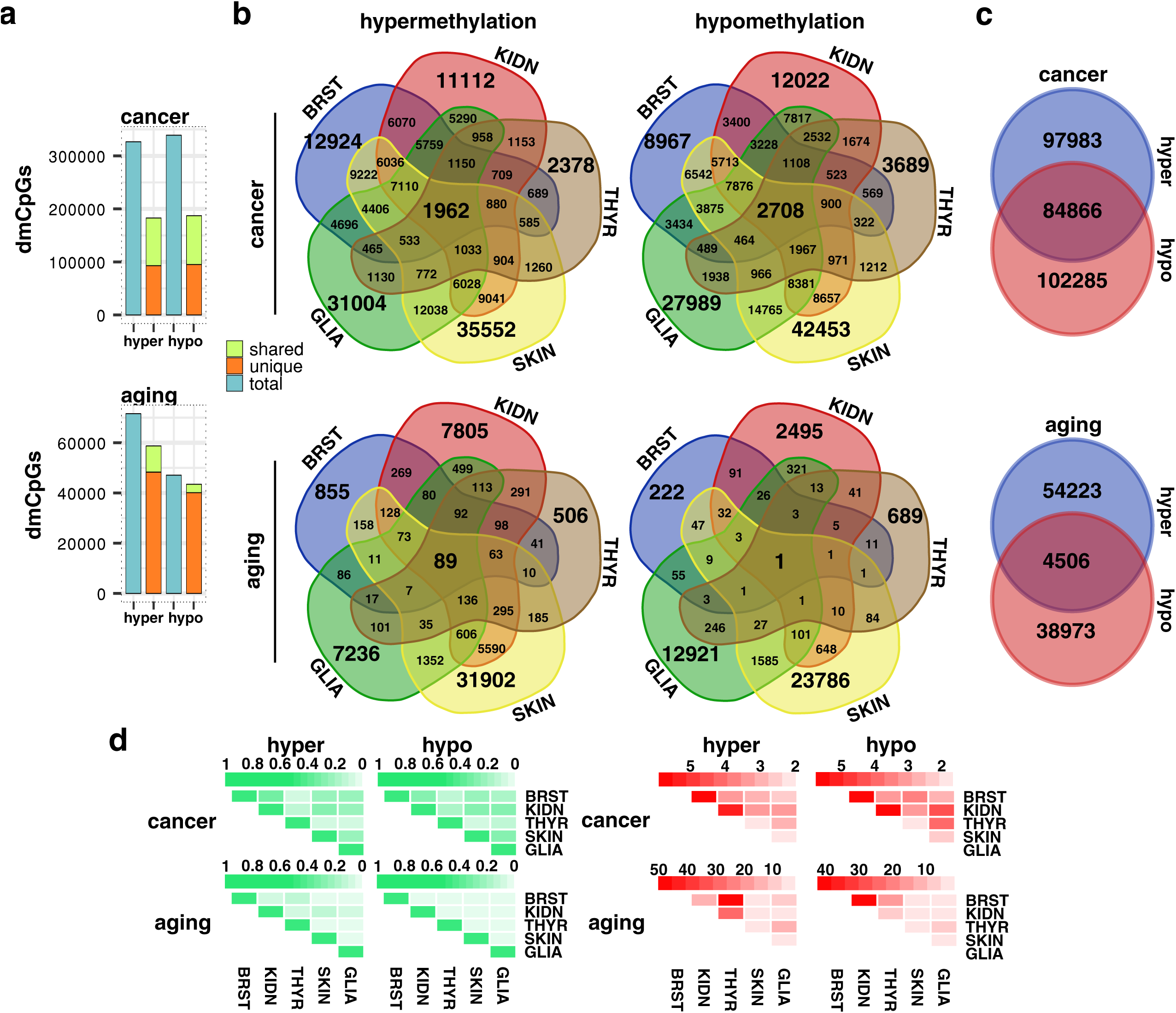
*DNA methylation signatures of aging and cancer. **a** Stacked barplots indicating, in blue, the total number of hyper- and hypomethylated dmCpGs detected between all of the tissues analysed in cancer and aging. Of these, the proportion of dmCpGs not shared between any tissues (unique, orange) or those shared by two or more tissues (shared, green) is shown in the adjacent barplot. **b** Venn diagrams depicting the number of differentially hyper- and hypomethylated CpGs in aging and cancer shared by the different tissues. **c** Venn diagrams showing the number and overlap of total non-redundant hyper- and hypomethylated dmCpGs detected in cancer and aging. **d** Heatmaps showing pairwise comparisons between sets of probes: in green, Jaccard Indices; in red, Odd ratios (all enrichment Fisher’s tests *P*<0.001).*

We also identified a subset of dmCpG sites in aging and cancer that could potentially be either hyper- or hypomethylated depending on the tissue type involved (Fig. 3c). Despite the substantial under-enrichment, this type of ambiguous dmCpG site was relatively more abundant in cancer than in aging (Fisher’s test *P*<0.001 for both, ORs = 0.65 and 0.56; expected hypergeometric means, EHMs = 94,610 and 7,060; and Jaccard Indices, JIs = 0.30 and 0.05, respectively). Interestingly, when examining dmCpGs shared by two or more tissues (Additional file 1: Figure S4b), this under-enrichment became more pronounced such that CpGs that were thus affected in more than one tissue were less likely to behave differently in other tissues.

### Similar chromatin signatures of DNA hypermethylation in aging and cancer

To identify possible chromatin marks associated with hypermethylated CpG sites in aging and cancer, we compared the hypermethylated CpG sites identified in this study with previously published ENCODE and NIH Roadmap Epigenomics ChIP-seq data on the histone modifications H3K4me1, H3K4me3, H3K27ac, H3K36me3, H3K27me3, H3K9me3 across 98 different cell and tissue-types (see Methods). The results confirmed an enrichment of hypermethylated CpG sites in repressive histone modifications H3K27me3 and H3K9me3 and active histone modifications H3K4me1 and H3K4me3 in both in aging and cancer (Fig. 4a, upper panel; Additional file 9: Table S8).

**Fig. 4.**
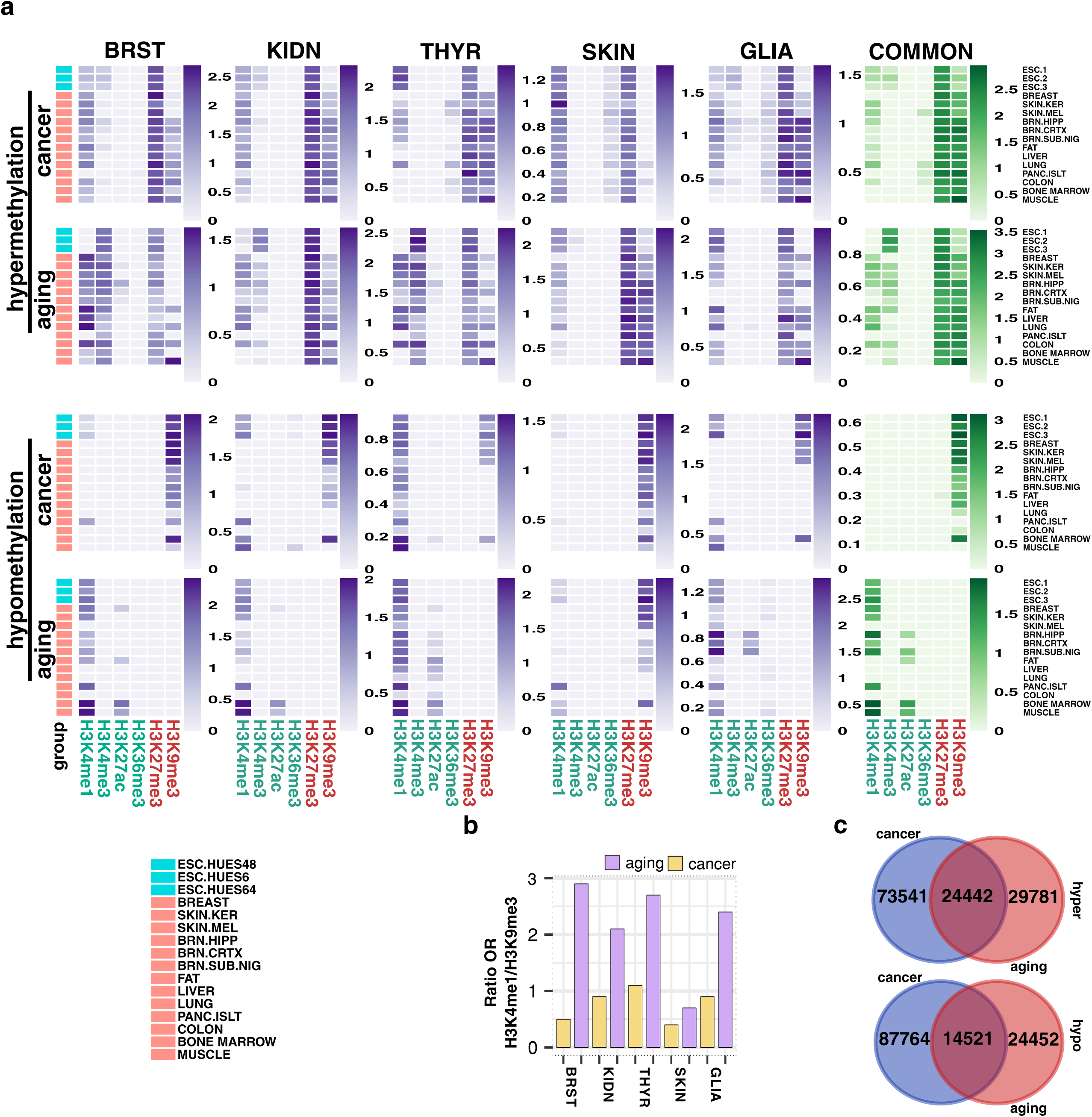
*Distinct chromatin signatures of hyper- and hypomethylation in aging and cancer. **a** Heatmaps depicting significant (*P*<0.05) over-enrichment of hyper- and hypomethylated dmCpG sites with different histone marks in aging and cancer, in a selection of 16 cell- and tissue-types (see Additional file 9: Table S8 for 98 full cell and tissue-types). Color code indicates the significant enrichment based on log2 odds ratio (OR). Common signatures are calculated from hyper- and hypomethylated dmCpGs shared between 5 tissues for cancer (1,962 and 2,708 probes, respectively) or 3 tissues for aging (904 and 106 probes, respectively) (see Additional file 7: Table S6 for CpG lists). **b** Barplots indicating the ratio of OR for the H3K4me1/H3K9me3 marks associated to hypomethylated dmCpGs in aging and cancer. The ratio is calculated by taking the mean of OR of those tracks with significant over-enrichment for each histone mark and dividing the obtained numbers. **c** Venn diagrams showing the number and overlap of total non-redundant hyper- and hypomethylated dmCpGs detected in cancer and aging. dmCpGs that were only hypermethylated or only hypomethylated between all tissues were chosen for the comparison.*

Notably, these similarities became more pronounced when examining dmCpGs shared by all five tissue types in cancer, or three of the tissue types in aging (low numbers of common probes, due to tissue-specificity of aging dmCpGs, hindered analysis of dmCpGs shared by more tissues). All the tissue-types analysed were strongly enriched in the repressive histone modification H3K27me3 in both aging and cancer. In contrast, enrichment in H3K9me3 was more pronounced in thyroid and brain tumours, as well as aged skin (Fig. 4a, upper panel). The enrichment pattern of active histone modifications was also dependent on tissue type. In cancer, H3K4me1 was more abundant than H3K4me3 in both embryonic stem and adult cells/tissues, and this was also observed in aged skin and brain samples. However, aging of breast, kidney and thyroid tissue was primarily associated with H3K4me3 in embryonic stem cells and with H3K4me1 in adult tissues/cells (Fig. 4a, upper panel). Interestingly, the embryonic stem cell signature was comparable to other tissue signatures, although, when present, the H3K4me3 mark was more evident in embryonic stem cells, especially in the aging context. Collectively, these results suggest that chromatin signatures of DNA hypermethylation are similar in aging and cancer.

### Distinct chromatin signatures of DNA hypomethylation in aging and cancer

To determine whether the chromatin signatures of DNA hypomethylation were also similar in aging and cancer, we compared the hypomethylated CpG sites identified in our study with data from the same histone modifications as described in the earlier analyses. Interestingly, the results showed that hypomethylated CpG sites in cancer were enriched in the repressive H3K9me3 histone modification, whilst in aging, hypomethylated CpGs were more enriched in the activating histone mark H3K4me1 (Fig. 4a, lower panel; Additional file 9: Table S8). There were though exceptions to this general trend: hypomethylated CpG sites in thyroid tumors were also enriched at H3K4me1, and hypomethylated DNA sequences in aged skin were mainly co-associated with H3K9me3-marked DNA regions. Nevertheless, the ratio H3K4me1/H3K9me3 was always higher in aging than in cancer (Fig. 4b). Moreover, when analysing the dmCpGs shared by all five tissues in cancer or at least three tissues in aging, these distinct chromatin signatures became much more evident. In sum, these results indicate that, in contrast to DNA hypermethylation, chromatin signatures of DNA hypomethylation in aging and cancer differ considerably.

After deriving the chromatin signatures we performed validation analyses on two additional datasets: the first related to tissue from TCGA control and lung adenocarcinoma, and the second to whole blood from the classical Hannum et al. dataset [24](Additional file 1: Figure S5; see Additional file 10: Table S9 for additional information). Interestingly, we were unable to find aging-related methylation changes in normal lung tissue using our pipeline. The magnitude and distribution of the hyper- and hypomethylation changes in lung cancer and whole blood aging followed the same trends as observed for the other datasets (Additional file 1: Figure S5a, see Additional file 3: Table S2 for list of dmCpGs). The histone enrichment analyses revealed the same hypermethylation signature previously found for cancer and aging, and very clear and different hypomethylation signatures of H3K9me3 for lung cancer and H3K4me1/3 for whole blood aging (Additional file 1: Figure S5b, Additional file 9: Table S8).

To identify possible explanatory factors underlying the different chromatin signatures of DNA hypomethylation in aging and cancer, we compared the overlap between either hypermethylated or hypomethylated CpGs across tumors and their corresponding age-related tissues (Fig. 4c). This approach revealed that the common overlap between hypermethylated CpGs (24,442 CpGs) was higher than expected by chance (Fisher’s test *P*<0.001, OR = 2.61; EHM = 14,689; JI = 0.20) (Fig. 4c, upper panel). However, despite the overlap between hypomethylated CpGs (14,521) also being slightly higher than expected (Fisher’s test *P*<0.001, OR = 1.60; EHM = 11,021; JI = 0.12) (Fig. 4c, lower panel), the overall trend observed in this case was weaker than for the hypermethylated CpGs. Furthermore, most of the hypomethylated probes shared by cancer and aging belonged to skin dmCpGs, providing evidence for its similar cancer and aging hypomethylation signatures. When skin tissue was discarded from the analysis (Additional file 1: Figure S6) the observed over-enrichment disappeared in the case of DNA hypomethylated probes, although it remained in the hypermethylation scenario (Fisher’s tests, both *P*<0.001, ORs = 0.8 and 3.0; EHMs = 5,357 and 7,568; JIs = 0.04 and 0.12, respectively), reinforcing the observed overlapping differences between hyper- and hypomethylated probes.

### Functional characterization of differentially methylated sites in aging and cancer

To determine the possible functional consequences and genomic coincidence of the different histone marks of DNA hyper- and hypomethylation in aging and cancer, we performed an enrichment analysis of NIH Roadmap and ENCODE Hidden Markov Model (HMM) defined “chromatin states” across the same 98 human cell and tissue types used in the previous analyses (see Methods). In total, eighteen states were used for the segmentation of the genome, which were then grouped to highlight predicted functional elements.

As suggested by the earlier chromatin signature analyses, hypermethylated CpGs in both aging and cancer were enriched in states associated with bivalent chromatin domains (i.e. those formed by the combination of repressive histone mark H3K27me3 and activating histone marks H3K4me3/1), polycomb repressive domains and repeat/ZNF genes. These patterns became more evident when examining the dmCpGs shared by all five cancer tissues or at least three aging tissues (Fig. 5a; see Additional file 1: Figure S7 for tissue-specific signatures and Additional file 11: Table S10 for full data in all 98 cell- and tissue types). Hypomethylated CpG sites in cancer were enriched in chromatin states associated with heterochromatin and repeat/ZNF gene domains and, to a lesser extent, polycomb repressive domains. In contrast, DNA hypomethylation in aging was primarily associated with chromatin states related to DNA enhancers. Again, these marks were more pronounced in aging dmCpGs shared by at least three tissues. As occurred with chromatin signatures, hypomethylation chromatin state differences were weaker in skin and thyroid, albeit that the ratio of change of chromatin states always followed the same behavior (data not shown). Collectively, these results support the notion that DNA hypomethylation might have a different functional role in aging as compared to cancer.

**Fig. 5.**
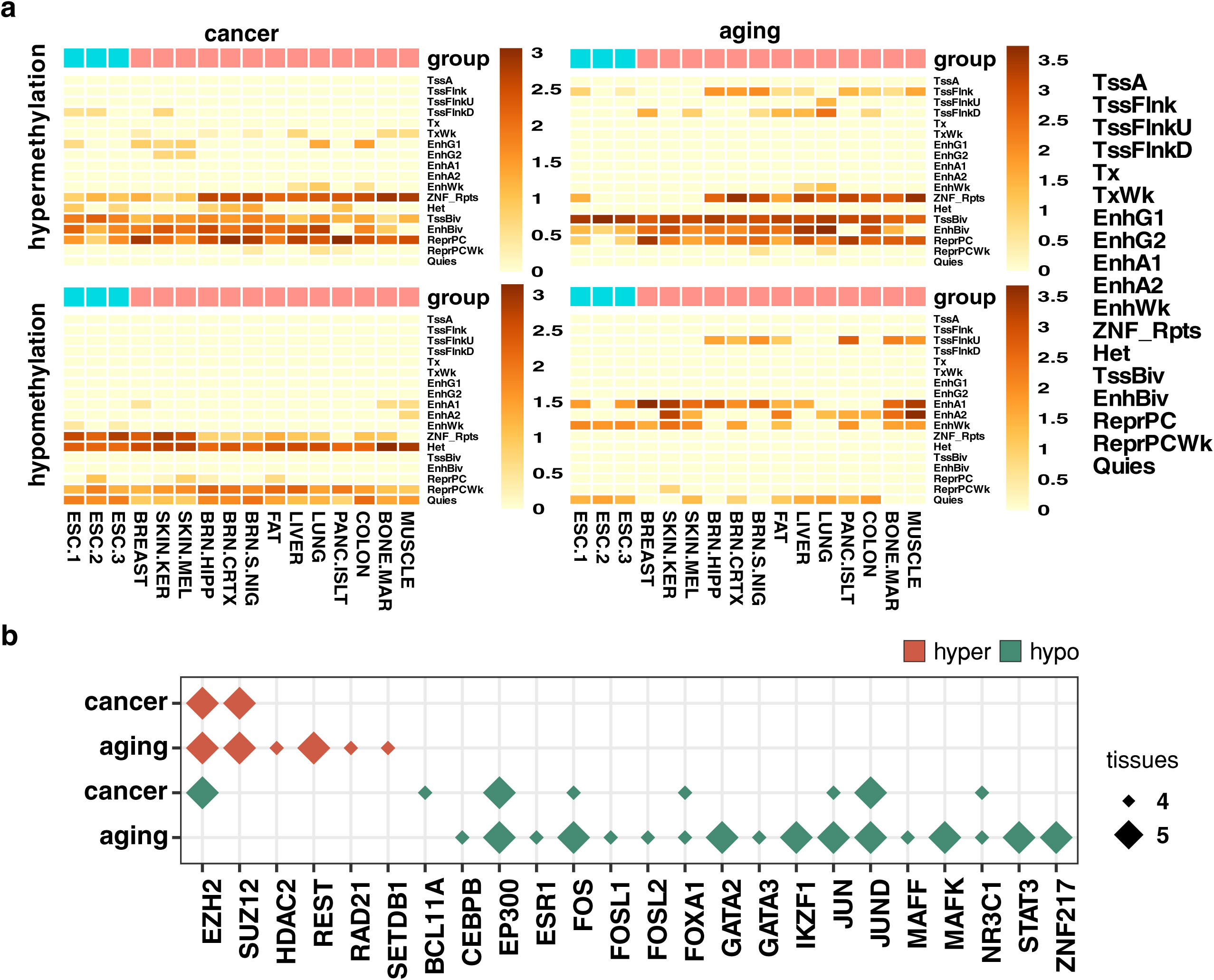
*Differential chromatin-state signatures of hyper- and hypomethylation in aging and cancer. **a** Heatmaps displaying significant (*P*<0.05) over-enrichment of hyper- and hypomethylated dmCpG sites with different chromatin states in aging and cancer, in a selection of 16 cell- and tissue types (see Additional file 1: Figure S7 for tissue-specific signatures, and Additional file 11: Table S10 for 98 full cell- and tissue-types). Color code indicates the significant enrichment based on log2 odds ratio (OR). Common signatures are calculated from hyper- and hypomethylated dmCpGs shared between 5 tissues for cancer (1,962 and 2,708 probes, respectively) or 3 tissues for aging (904 and 106 probes, respectively) (see Additional file 7: Table S6 for CpG lists). **b** Panel indicating significant (*P*<0.05) transcription factor enrichment at hyper- and hypomethylated dmCpG sites in aging and cancer that occurred in at least 4 or 5 tissues (see Additional file 1: Figure S8 for tissue-specific results and Additional file 12: Table S11 for full data). Only the most representative transcription factors (those that appeared as significantly over-enriched in at least 3 tracks or with an OR>3 in any track) were selected for data representation.*

In order to increase our understanding of the functional context of the chromatin signatures characterized, we compared the dmCpG sites identified in this study with publicly available ENCODE ChIP-seq data on transcription factor binding sites in 689 datasets corresponding to 188 transcription factors across 91 different cell types (see Methods) (Fig. 5b; see Additional file 1: Figure S8 and Additional file 12: Table S11 for full tissue-specific results). As expected, hypermethylated CpGs in aging and cancer were associated in all tissues with the presence of EZH2 and SUZ12, components of the polycomb complex which directly deposits the H3K27me3 mark. Interestingly, aging hypermethylation dmCpGs were specifically associated with other types of transcription factors in various tissues, such as REST, HDAC2, RAD21, and SETDB1. Transcription factor enrichment at hypomethylated dmCpG sites was more heterogeneous than, but different from that of hypermethylated sites. Enrichment of similar factors was found for cancer and aging, for example, EP300, FOS and JUN, among others. As observed before, specific aging enrichment was found, such as GATA2/3. In this case, aging hypomethylated dmCpG sites tended to display a more marked enrichment of most of the cancer hypomethylation factors and, additionally, revealed the presence of other family- or function related proteins, like FOSL1/2, MAFF, MAFK and STAT3. When examining enrichment at common dmCpG sites shared by different tissues in cancer and aging, the initial observations were further confirmed (Additional file 1: Figure S9).

We also performed gene and KEGG pathway ontology analyses (Fig. 6a; Additional file 13: Table S12; see Additional file 1: Figure S10 for tissue-specific results). Hypermethylated CpGs in both cancer and aging belonged to genes that were mainly related to developmental functions. While genes containing hypomethylated CpGs in cancer were associated with extracellular signaling, those for aging were, in general, much less enriched in any gene ontology. In the case of KEGG pathways (Fig. 6a; Additional file 13: Table S12 and Additional file 1: Figure S10), hypermethylated CpGs in both cancer and aging shared enrichment for several ontologies, many related to cell metabolic and signaling pathways. In this respect, hypomethylated CpGs in cancer had some ontologies in common while others were specific. Once again, aging hypomethylated CpGs exhibited much less enrichment in any function.

**Fig. 6.**
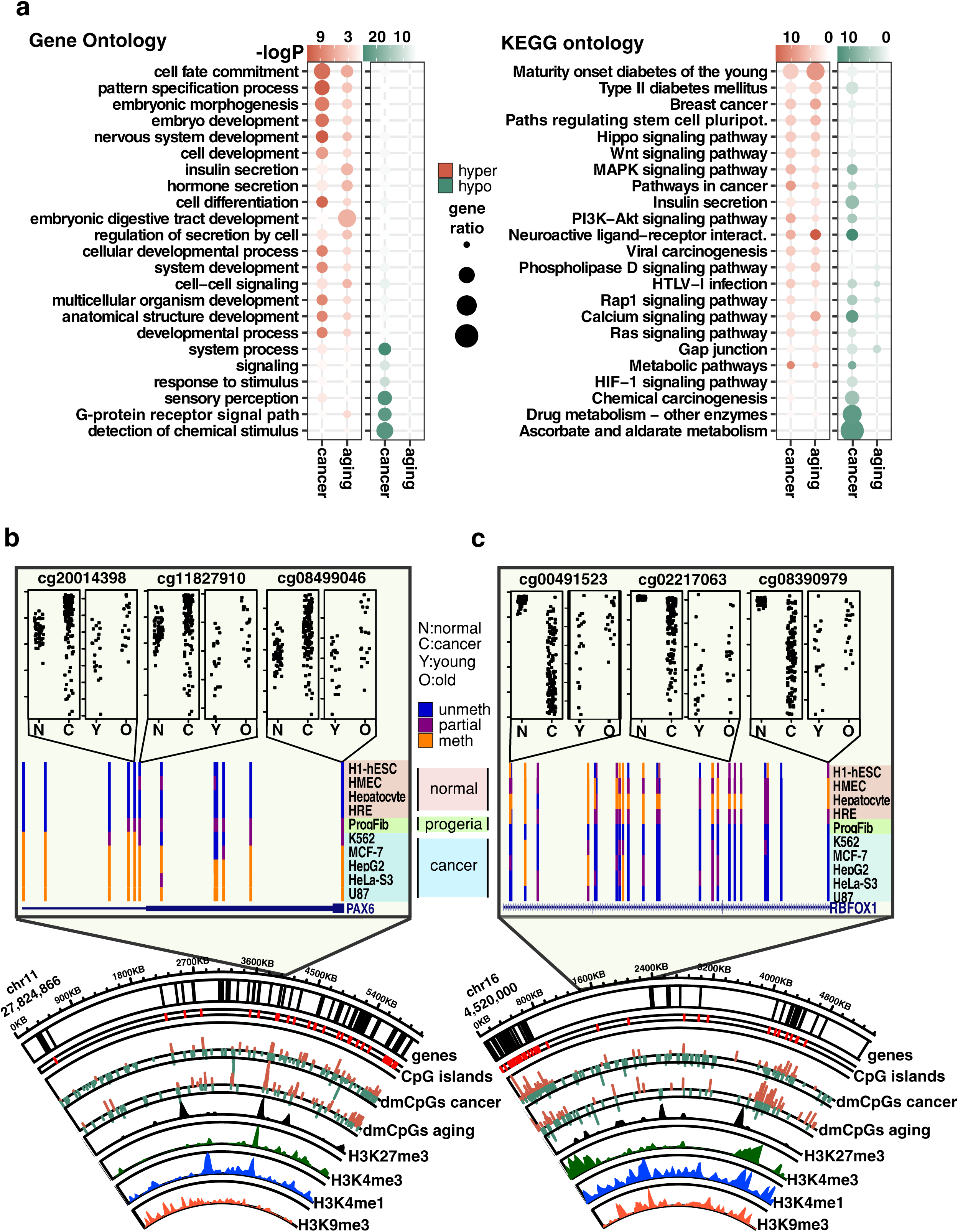
*Dissimilar functional context of differentially methylated CpGs in aging and cancer. **a** Panels indicating gene and KEGG pathway ontology enrichment for common hyper- and hypomethylated dmCpGs shared between 5 tissues for cancer (1,962 and 2,708 probes, respectively) or 3 tissues for aging (904 and 106 probes, respectively) (see Additional file 7: Table S6 for CpG lists and Additional file 1: Figure S9). Color code indicates the significance of the over-enrichment based on log10 P-value. Size of circles indicates gene ratio, calculated as the ratio of number of identified hits with respect to the total number of components in a given ontology. **b** and **c** Circular representation of illustrative genomic locations indicating hyper- (red) and hypomethylated (green) dmCpG sites in cancer and aging. Inner tracks display chromatin marks (H3K27me3, H3K4me3, H3K4me1 and H3K9me3, respectively), corresponding to ChIP-seq peak data from ESC E014 NIH Roadmap track (circos, lower panel). Two examples of hypo- and hypermethylated genes are highlighted from the circular figures, displaying methylation data obtained from ENCODE/HAIB methyl450 track set from UCSC Genome Browser (hg19) and plots of methylation values extracted from a representative example of our aging and cancer analyses corresponding to glia tissue (upper panel, depicted CpGs also displayed a similar trend in kidney and skin analyses, data not shown).*

In order to exemplify the similarities and disparities observed for DNA methylation in aging and cancer, we focused on a number of significant dmCpGs from two particular genomic regions, located in chromosomes 11 and 16. (Fig. 6b and 6c). We observed a substantial correlation between bivalent posttranslational histone modifications, especially H3K27me3 and H3K4me1/3, and presence of hypermethylated probes in aging and cancer. On the other hand, DNA hypomethylated regions were more frequently located near H3K9me3 or H3K4me1 peaks (bottom panel Fig. 6b and 6c) as outlined in our previous histone enrichment analyses. A detailed inspection of the common genes with most abundant dmCpGs in aging and cancer revealed a similar trend towards DNA hypermethylation at the boundaries of the gene *PAX6* (Fig. 6b, top panel). Interestingly, a representative set of cancer cell lines, as well as fibroblasts derived from patients with Hutchinson-Gilford progeria, also display higher levels of DNA methylation when compared to normal cells in these differentially methylated regions (Fig. 6b, middle panel). On the contrary, the abovementioned pattern was mainly reversed in the case of the *RBFOX1* gene, located in a region which was preferentially hypomethylated in cancer (Fig. 6c, top and middle panel).

### Correlations between CpG methylation and gene expression in aging and cancer

Despite the similarities and disparities observed here for the chromatin signatures of aging and cancer, the functional impact of these methylation changes in the control of gene expression remained unknown. To address this issue, we focused on kidney tissue (KIRC) as this TCGA dataset displayed a reasonable number of control and cancer patients with paired methylation and gene expression data. We initially performed differential gene expression analyses comparing young versus old kidney tissue or normal versus tumoral kidney samples (Fig. 7a, Additional file 14: Table S13). These results allowed us to identify a total of 13 and 20,678 differentially expressed genes (DEGs) in aging and cancer conditions, respectively. The majority of the aging DEGs were also found in cancer, including, for example, the *CKM* gene, which contained a dmCpG in the proximity of its promoter (Fig. 7b), was differentially expressed in both processes (Fig. 7c) and displayed a considerable negative correlation between DNA methylation and gene expression in normal kidney (Spearman correlation = −0.37, Fig. 7d). To further explore the potential relationships between CpG methylation and gene expression in these processes, and due to the reduced number of DEGs observed in the aging context, we decided to perform all potential pairwise correlations between DNA methylation and gene expression using cancer or aging related dmCpGs and genes expressed in a subset of normal kidney tissue samples (n = 18). This approach enabled us to quantify the extent to which CpGs whose methylation status changes in cancer and aging originally influence gene expression in normal tissue.

**Fig. 7.**
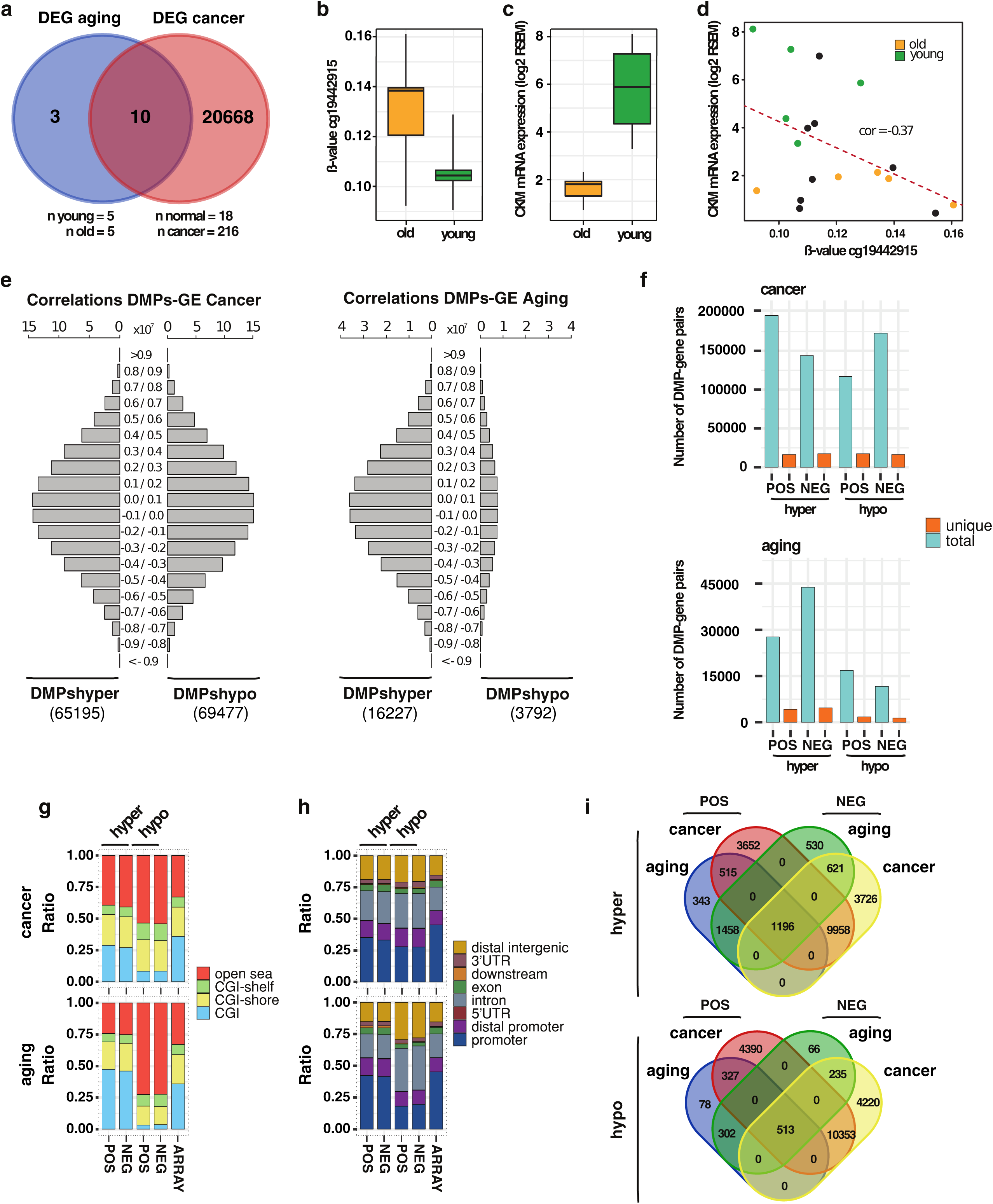
*Relationships between DNA methylation and gene expression in aging and cancer. **a** Venn diagrams illustrating the overlap between DEGs in aging and cancer in the KIRC dataset (see Additional file 14: Table S13 for DEG lists). **b** Boxplot depicting the DNA methylation β-values of the CpG cg19442915 in old and young individuals (n = 5) from the KIRC aging condition. **c** Boxplot showing the gene expression values (RSEM) of the CKM gene in old and young individuals (n = 5) from the aging condition of the KIRC dataset. **d** Scatterplot showing the Spearman correlation between DNA methylation (cg19442915) and gene expression (CKM gene) in 18 normal kidney samples. Colored dots indicate old or young individuals used for the aforementioned aging comparisons. **e** Histograms representing the number of pairwise correlations that are contained in a given correlation window (from 1 to −1) obtained as the result of computing the correlation between β-values of dmCpGs identified in cancer or aging and gene expression levels (RSEM) of genes expressed in the KIRC dataset. The number of dmCpGs used for each of the comparisons is indicated at the bottom. **f** Barplots depicting the number of total (blue) and unique (orange) dmCpG-gene pairs identified in the previous analysis which displayed correlation scores above 0.9 (pos) or below −0.9 (neg) in cancer (top) or in aging (bottom) conditions. **g** Stacked barplots indicating relative distribution of unique dmCpGs obtained from the previous correlations according to their CpG island status. **h** Stacked barplots indicating relative distribution of unique dmCpGs obtained from the previous correlations according to their gene location status. **i** Venn diagrams illustrating the overlap between dmCpGs identified in aging and in cancer which displayed strong positive or negative correlations (>0.9 or < -0.9) with genes expressed in the normal kidney dataset.*

We computed a total of 2.58e^09^ and 3.84e^08^ correlations between cancer and aging related dmCpGs, respectively, and genes expressed in the normal KIRC dataset (Fig. 7e). Despite the considerable difference in number of dmCpGs between cancer and aging, when compared to the total possible number of correlations, we found similar percentages of strong correlations between DNA methylation and gene expression in both processes (Additional file 15: Table S14). Moreover, these proportions were also higher than those observed when sampling random probes from the array and computing their correlations (see Additional file 1: Figure S11). These results indicate that both cancer and aging related dmCpGs are enriched in CpGs that can influence, to some extent, gene expression in kidney tissue.

A more detailed inspection of the strongest correlations (>= 0.9 or <= −0.9) identified in these datasets revealed that, in cancer, the number of unique dmCpG-gene pairs remained similar (~20,000) regardless of the direction of the observed correlation (Fig. 7f, top). Furthermore, while the number of unique dmCpG-gene pairs in the aging context was much reduced (~3,000), the proportion compared to the total number of strong correlations observed in a given dataset remained, to a great extent, similar (Fig. 7f, bottom). Interestingly, the differences between the genomic distributions of the unique hyper- and hypomethylated -dmCpG-gene pairs identified in aging or in cancer followed the same trend to those observed for the genomic distribution of the hyper- and hypomethylated -dmCpGs identified in each of these processes (Fig. 2b-c), with hypermethylated dmCpGs being more enriched in CpG islands in aging as compared to the array (Fisher’s tests; both *P*<0.001, ORs = 2.0, 1.4, for positively and negatively correlated dmCpGs, respectively), in contrast to the enrichment at open-sea locations (Fisher’s tests; all *P*<0.001, ORs = 4.6, 4.1, 2.3 and 2.5 for positively and negatively correlated dmCpGs in aging and cancer, respectively) and intronic regions (Fisher’s tests *P*<0.002, <0.03, <0.001 and <0.001, ORs = 2.2, 1.9, 1.6 and 1.7 for positively and negatively correlated dmCpGs in aging and cancer, respectively) of the hypomethylated dmCpGs in both processes (Fig. 7g-h). It is worth noting that the distribution of the unique hypermethylated-dmCpGs which also control gene expression was more enriched in open sea locations as compared to dmCpGs in general in both aging and cancer (Fig. 7g as compared to Fig. 2b).

Finally, we compared the unique dmCpGs that displayed strong correlations between DNA methylation and gene expression in aging or in cancer (Fig. 7i). We observed an extensive overlap between probes that displayed positive or negative correlations in the two processes (Fisher’s tests, all *P*<0.001, ORs = 908, 2244, 205, and 171; JIs = 0.57, 0.54, 0.57 and 0.54 for aging and cancer hyper- and hypomethylated CpGs, respectively). This fact might explain their similar genomic distributions (Fig. 7g-h), indicating that most of these dmCpGs could play a dual role in the control of their different gene expression targets. We also observed a considerable overlap between dmCpGs associated to gene expression identified in aging and cancer processes (Fisher’s tests, both *P*<0.001, ORs = 5.1 and 9.1; JIs = 0.06 and 0.03, for hyper- and hypomethylated CpGs, respectively). Interestingly, regardless of whether the DNA methylation change was towards hyper- or hypomethylation, ~ 60-70% of the aging-related dmCpGs which controlled gene expression were also present in the group of cancer related dmCpGs (Fig. 7i). These results point towards similarities of cancer and aging related dmCpGs in the control of gene expression in normal tissue, despite the fact that the number of cancer related dmCpGs is clearly larger than their aging counterparts.

## Discussion

Although it is widely accepted that cancer is an age-dependent disease, the underlying molecular mechanisms are still poorly characterized. Interestingly, recent studies have proposed that DNA methylation might play an important role in the tumorigenic process because genes aberrantly hypermethylated in both aging and cancer are enriched in bivalent chromatin domains and polycomb bound regions in embryonic stem cells [4–6,9–11]. Loss of DNA methylation at specific DNA regions has also been independently described in aging and cancer [27]. However, it was unclear whether DNA hypomethylation also shared similar chromatin signatures in both processes, something which is conceptually relevant to understanding the role of 5 methylcytosine (5mC) as a possible molecular link between age decline and tumor development.

In agreement with previously published data [11,28], we observed that the number of aging- and cancer associated DNA methylation changes was variable and, in the case of aging, had a marked tissue-type dependent component. In general, cancer displayed bidirectional changes while, strikingly, hypermethylated CpG sites were predominantly observed for the aging process. These results, which are ostensibly in contrast with the classically described global hypomethylation changes in cancer and aging, might potentially arise from the limitations of our study. As the methylation arrays used in our analyses mainly interrogate genetic elements and do not include repeated DNA, which covers a substantial fraction of the genome and frequently loses DNA methylation in tumors and aged cells, the genome-wide landscape may be different [16,27]. Nonetheless, epigenetic signatures have been successfully derived previously using array technology [9–11]. Despite a careful normalization being applied for each of the comparisons, slight deviations in the number of dmCpGs in particular tissues could be influenced by the origin and type of the samples used in our statistical analyses, especially in the case of glia and skin controls, since the data processed were obtained from a number of different public databases. Furthermore, the changes in cell type composition that occur with age and cancer are also well known confounding factors that could affect our datasets [29]. However, the application of the SVA method of correction (and Houseman correction for blood) and the use of a pure-cell dataset such as the glia dataset allowed us to tackle this issue in two different ways [30]. Additionally, the use of the blood validation dataset [24] allowed us to verify the reliability of our workflow, as in terms of whole blood dmCpGs we obtained 89% concordance with previous studies using the same data [11].

To compare the behavior of the DNA methylation changes in aging and cancer we first analysed the genomic distribution of dmCpGs and, in line with previously published reports [31–34], we found that hypomethylated CpGs were enriched at open sea DNA regions, principally those that were intronic and intergenic, irrespective of the type of process. The distribution of hypermethylated CpGs was found to be similar to that of the array, which is to a certain extent to be expected because it was designed to interrogate a promoter- and CpG dense-biased portion of the genome. Nonetheless, hypermethylation changes always occurred in far more CpG-dense regions than hypomethylation changes [13,32–34]. Our study allowed us to ascertain that the observed effect was especially pronounced for aging dmCpGs. Very interestingly, while this polarization in genomic location of hyper- and hypomethylated CpGs is more evident with regards to the aging process, we found methylation changes to be much more pronounced in tumor samples. This observation is compatible with previous descriptions of epigenetic clock acceleration in cancer [26,35].

When studying the potential effect of tissue type on DNA methylation changes we found, in agreement with recently published data [36], that DNA methylation changes in different tumor types were surprisingly similar, regardless of the tendency of the alteration. This observation is conceptually relevant because it has classically been considered that different tumor types are characterized by specific DNA methylation signatures [37,38], especially in the case of DNA hypermethylation. In this sense, our data confirm that, although different tumor types might display specific DNA methylation patterns, there is a significant common nexus between them. The analysis of the DNA methylation changes observed with respect to the aging process also revealed significant overlap between tissue types. It is possible that these results are affected by the variability in the sizes of the sets of probes detected in the aging analysis, however, everything considered, in this work both cancer and aging manifest tissue-independent changes which could reflect core characteristics of each process.

Our data revealed that dmCpGs shared by two or more tissues were much less likely to have different behaviors in other tissues, perhaps pointing towards non-stochastic and possibly functional roles for these CpGs. Overall, there was still a large proportion of cancer hypermethylated CpGs in a certain tissue that were able to be detected as hypomethylated CpGs in other tumoral tissues, indicating that many of these CpGs have tissue-specific directional changes that could be explained in two possible ways: a heterogeneous epigenome state in which the affected sites suffer a stochastically directional drift, perhaps restricted by an initial configuration in the original tissue [39,40]; or a functionally directional drift in which different tumors acquire different epigenome-states that aid in cancer progression [41,42].

The systematic DNA methylation analyses described in this study confirm that DNA hypermethylation in aging and cancer is associated with the same set of histone marks, including the repressive H3K27me3 and H3K9me3 marks, and the activating H3K4me1/3 post-translational modifications. Chromatin-state analysis revealed that the hypermethylation-associated H3K27me3 and H3K4me1/3 marks configured bivalent chromatin domains, as has been extensively described in embryonic stem cells [4–6,9–11]. Moreover, our data reveal that this chromatin signature is not restricted to only embryonic stem cells, but rather this trend should be considered an extended, global tissue-independent chromatin signature of DNA hypermethylation in aging and cancer. Interestingly, Chen and colleagues [36] have recently demonstrated that normal-tissue signatures are better predictors of DNA hypermethylation changes than ESC signatures. Furthermore, hypermethylation changes were also associated with the repressive histone mark H3K9me3 [6,43], which was correlated to ZNF genes and DNA repeats in our chromatin state analyses, and which might have a potential relationship with the malignant transformation process [43,44].

Regarding DNA hypomethylation, our results showed that age-related DNA hypomethylation is associated with the activating histone posttranslational modification H3K4me1, which supports previously published data [11]. A slight tendency for enrichment of H3K27Ac, a histone mark characteristic of active enhancers [45], was also detected in our analyses. Intriguingly, the chromatin signature of DNA hypomethylation in cancer was substantially different, with most tumor types being primarily enriched in the posttranslational repressive histone modification H3K9me3, a relationship that has been investigated in colon and breast cancer [46,47]. This observation might be conceptually relevant because DNA methylation has been proposed to be a molecular link between aging and cancer [27,48,49]. However, our results suggest that the role of DNA methylation as a possible link between aging and cancer is more complex than previously proposed. Importantly, even though most of the observed DNA methylation changes in aging were tissue specific, we were able to describe a common chromatin signature characteristic of the aging process. This observation points towards the notion that there might be non-stochastic underlying molecular mechanisms, and that these differ between aging and cancer.

The different chromatin signatures of DNA hypomethylation in aging and cancer might have important functional consequences. This was clearly evidenced by the analysis of the chromatin states. Indeed, DNA hypomethylation in cancer was associated with heterochromatin DNA regions, which is in line with previous work [31,46]. It is curious that chromatin segmentation analysis of chromatin states containing H3K9me3 revealed differing functionality of this mark depending on whether it was associated with cancer and aging DNA hypermethylation (ZNF genes and repeats) or simply DNA cancer hypomethylation (heterochromatin).

In contrast, chromatin marks of DNA hypomethylation in aging were associated with enhancers, reinforcing previous observations performed with the Infinium HumanMethylation27K Beadchip platform [13]. As DNA methylation changes in enhancers have been shown to play an important role in gene regulation [50–52], our results suggest that DNA hypomethylation during aging might have a different functional role in gene regulation compared to DNA hypomethylation changes in cancer.

With regards to the potential effectors of the distinct chromatin signatures, enrichment analyses of transcription factors revealed the presence of EZH2 and SUZ12 polycomb components at DNA hypermethylated sites, both in cancer and aging. Specific aging hypermethylation-associated factors were also observed in our comparisons. As reported in previously published data in blood [34], aging hypermethylation was associated in many tissues with REST, which has been shown to participate in DNA demethylation in mouse [53] and has also been correlated with longevity [54]. Thus, its age-related loss might lead to the observed DNA hypermethylation at its binding sites. Other factors, such as SETDB1, which establishes the H3K9me3 mark [55], were also related to aging but not cancer hypermethylation.

Concerning DNA hypomethylation, transcription factors such as FOS, JUN and JUND were detected at both cancer and aging hypomethylated CpG sites, but again aging displayed stronger and more varied enrichment, and included the presence FOSL1/2. Interestingly, these factors, which form the dimer AP-1, have recently been described to locate to active enhancer genomic regions [56]. Other bZIP-domain factors like MAFF and MAFK were also related to aging hypomethylation. Additionally, STAT3, which has been associated with recruitment of the H3K4 methyltransferase SET9 at promoters [57], and GATA2/3 were also associated with aging hypomethylation sites. Altogether, these observations would imply that hypomethylation in aging displays a more marked functional context than that of cancer, exhibiting an increased enrichment of some factors also detected at cancer hypomethylated sites and other specific factors not found associated with tumoral changes.

Interestingly, our gene ontology analyses revealed similar gene functionalities affected by cancer and aging DNA hypermethylation, mainly related to developmental processes, which is in line with the methylation of bivalent chromatin promoters of developmental regulators in cancer and aging [7,9]. On the other hand, DNA hypomethylation in cancer was mainly associated with functions identified with cellular signaling, and much lower enrichments in gene functions were found for hypomethylated CpGs in aging. A preponderance of non-genic enhancer hypomethylation in aging could potentially explain this absence of gene function association in our data.

To date, the potential relationships between DNA methylation and gene expression have only been systematically analysed in a small subset of studies [58–60], and the potential effects of these relationships on aging and cancer are yet to be elucidated. To this end, we explored the establishment of potential correlations between these two processes using the TCGA KIRC dataset. While most correlative studies focus on CpGs located at particular genomic regions, such as DNA promoters [61] and cis-related correlations with the gene of interest [60], we performed a non-biased approach focusing on all the potential pairwise comparisons that could be identified between any significant dmCpG and the genes expressed in the context of normal kidney tissue. The limitations of these analyses did not allow us to distinguish between direct (i.e. mediated by the effects of the DNA methylation process itself) or indirect regulation of gene expression governed by the subsequent expression of other regulatory factors. Nonetheless, we observed that both aging and cancer dmCpGs influence gene expression to a similar extent, as these processes show the same proportions of strong correlations between DNA methylation and gene expression in normal kidney tissue. Moreover, we observed a similar number of positive and negative correlations between DNA methylation and gene expression, as described in Gutierrez-Arcelus et al. 2013, with most of these positively and negatively correlated dmCpGs overlapping substantially, suggesting that these CpG sites may play a dual role in the control of gene expression, or the involvement of other factors.

Finally, we found that most of the tumor types analysed in this study did not show age-associated DNA methylation changes, which is in agreement with the reprogramming of the epigenetic clock in cancer cells [26] and further supports the notion that malignant transformation amends the epigenetic alterations accumulated during aging. As an exception, we identified age-associated dmCpGs in thyroid tumors. Uncommonly, thyroid cancer includes age as a prognostic indicator in most staging systems [62], implying that these cancers do suffer age-related changes in their behaviour, a consequence of which could be our observed cancer-aging methylation changes. Intriguingly, thyroid tissue displayed the lowest level of DNA methylation changes in cancer and one of the lowest in aging. Although the reasons for the different behavior of DNA methylation changes in thyroid are currently unknown, they could be related to the good prognosis that typically characterizes this type of tumor. In fact, Yang Z. and collaborator’s “epiTOC” mitotic clock [35] shows thyroid cancer to have the least deviation from the behavior of its normal tissue. In this regard, future research should be conducted to address this issue.

## Conclusions

Our results indicate that hyper- and hypomethylated changes in aging and cancer each have similar genomic distributions and manifest tissue-independent trends in both processes. We confirm that chromatin signatures of DNA hypermethylation in aging and cancer are similar but, strikingly, we demonstrate that they are different for DNA hypomethylation. Collectively, our data suggest that the possible role of DNA methylation as a molecular link between aging and cancer is more complex than previously thought.

## Methods

### Data acquisition

All HumanMethylation450 BeadChip datasets used in this study are publicly available. DNA methylation data (Level 3) and clinicopathological features corresponding to normal or primary tumors from breast (BRCA), kidney (KIRC), thyroid (THCA), skin (SKCM) and glia (GBM) samples were obtained from TCGA consortium via UCSC Xena Public Data Hub (http://xena.ucsc.edu/) and NIH GDC Data Portal (https://portal.gdc.cancer.gov/). In order to increase the number of control cases in kidney, skin and glia tissues, these datasets were enlarged using additional samples from KIRP (TCGA), skin [25] and glia [23] respectively.

Tissues were chosen based on disease prevalence, control data availability and quality, and previous literature analyses in order to include both novel and pre-analysed tissues. We also performed analyses on two supplementary datasets: lung adenocarcinoma (LUAD) and control data from the TCGA consortium, and whole blood from a healthy cohort [24]. Extended information about control and tumor samples for each tissue type is shown in Table 1, Additional file 2: Table S1 and Additional file 10: Table S9.

### Data preprocessing

HumanMethylation450 BeadChip files containing β-values for ~450K probes had already been preprocessed and released by TCGA according to Data Level 3 guidelines. Probes located in chromosomes X and Y, probes overlapping genetic variants (SNP137Common track from UCSC genome browser), crossreactive and multimapping probes [63], as well as probes masked as NA (‘Not Available’) were discarded for downstream analyses. For each of the datasets, all samples (normal and tumor cases) were normalized using the BMIQ method [64] implemented in the R/Bioconductor package *ChAMP* (version 2.8.1)[65]. In the case of skin and glia tissues, since controls were obtained from various different studies, we performed a prior quantile normalization on these datasets [66], as implemented in the R/Bioconductor package *wateRmelon* (version 1.18.0) [67]. In the case of blood tissue, a specific cell-heterogeneity correction was applied as described by [68]. M-values were calculated from the normalized β-values through a logit transformation (R/Bioconductor package *lumi,* version 2.28.0)[69] and employed for downstream statistical analyses. Multidimensional scaling (MDS) and principal component regression analyses were used to identify potential confounding variables (R/Bioconductor package *Enmix,* version 1.12.3)[70]. To further correct for batch effects and unwanted sources of variation in the data, we performed a surrogate variable analysis [71] with the R/Bioconductor package *sva* (version 3.24.0). Surrogate variables for the different tissue datasets corresponding to “aging” (control cases or primary tumors) or “cancer” (control cases versus primary tumors) analyses were calculated employing the variable *age* or *sample_type* as the outcome of interest respectively. The number of latent factors for each of the different datasets was estimated using the “leek” method.

### Differential DNA methylation analyses

Differentially methylated probes (dmCpGs) in aging and cancer were calculated with the R/Bioconductor package *limma* (version 3.32.2)[72]. Briefly, a linear model between methylation levels as response variable, the variable of interest (either *age* group or *sample_type)* and the aforementioned surrogate variables was fitted for each of the analyses. For the calculation of age related dmCpGs, samples (control or tumor) were divided into quantiles in such a way as to obtain groups with sizes of n=15-30, and comparisons were performed between the upper (*OLD*) and the lower (*YOUNG*) quantile. Cancer related dmCpGs were calculated between normal tissue (*Solid Tissue Normal*) and tumor samples (*Primary Tumor*) as indicated in Table 1, Additional file 2: Table S1 and Additional file 3: Table S2.

The set of p-values obtained from the tests was adjusted for multiple comparisons using the Benjamini-Hochberg method to control for false discovery rate (FDR < 0.05). An additional threshold of shift size was applied, filtering out significant probes with M-value changes of less than 0.5, as has been suggested elsewhere [73].

Venn diagram representations of all possible logical relationships between statistically significant dmCpGs were generated with the online resource provided by the UGent/VIB bioinformatics unit (available at http://bioinformatics.psb.ugent.be/webtools/Venn/). Further enrichment analyses were performed by means of two-sided Fisher’s tests (*P*<0.05 significance threshold), measuring either effect size by odds ratio (OR), or the difference between observed counts and expected hypergeometric means, employing appropriate backgrounds of interrogated probes for the given context.

### Density of CpG analysis

For each of the probes in the HumanMethylation450 microarray, density of CpG was measured as the number of CpGs present divided by the number of those possible in a 2 kbp window centered on the CpG under study. Wilcoxon non-parametric tests were used to determine whether there were significant difference between the density distributions of the CpGs belonging to each subset of interest and the densities of the array probes in the background. A significance level of 0.05 was employed for all tests. Shift size was measured using median differences and Cliff’s Delta (D) (see Additional file 5: Table S4).

### CGI status and genomic region analysis

CGI island membership was assigned to each probe using Illumina’s 450K annotation with the R/Bioconductor package *IlluminaHumanMethylation450kanno.ilmn12.hg19* (version 0.6.0). Genomic region position was assigned using the R/Bioconductor packages *TxDb.Hsapiens.UCSC.hg19.knownGene* (version 3.2.2) and *ChIPseeker* (version 1.12.0)[74]. Statistical significance with respect to concrete CGI status or genomic regions was determined by two-sided Fisher’s tests (significance level *P*<0.05), and ORs were used as a measure of the association effect with respect to a particular feature (see Additional file 6: Table S5). Appropriate backgrounds which included all the probes interrogated by the HumanMethylation450 BeadChip array in each of the comparisons were used for statistical purposes.

### Region set enrichment analysis

Enrichment analyses were performed with the R/Bioconductor package *LOLA* (version 1. 4.0)[75], which looks for over-enrichment by conducting one-sided Fisher’s tests (*P*<0.05 significance threshold), by comparing overlap of probes (10 bp probe-centered windows) with the dataset of interest. Enrichment of histone marks was determined using histone ChIP-seq peak tracks (H3K4me1, H3K4me3, H3K27me3, H3K36me3, H3K9me3 and H3K27ac marks) from 98 epigenomes (primary tissues, cultures and cell lines) obtained from the NIH Roadmap and ENCODE projects [76,77] and integrated in the *LOLA* extended software (datasets obtained from http://databio.org/regiondb) (see Additional file 9: Table S8). The same method was employed for chromatin-segment analysis using NIH Roadmap’s ChromHMM expanded 18-state model tracks for the same 98 epigenomes (see Additional file 1 Figure S7 and Additional file 11: Table S10, custom database generated with data obtained from http://egg2.wustl.edu/roadmap/). In a similar fashion, ChIP-seq peak tracks from ENCODE for transcription factor binding sites (TFBS) comprising 689 datasets corresponding to 188 TFs analysed in 91 cell and tissue types were employed for TFBS enrichment analysis (datasets integrated in *LOLA* core region database, http://databio.org/regiondb, see Additional file 12: Table S11).

### DNA methylation age analyses

DNA methylation age of normal and tumoral samples was predicted by applying Horvath’s age predictor as implemented in R/Bioconductor package *wateRmelon* (version 1.18.0)[26,67]. The correlation of the predicted age thus obtained with chronological age was measured with the Pearson product-moment correlation coefficient using R package *stats* (version 3.3.3), which measures significance using a Student’s-t sampling distribution.

### Gene and KEGG ontology analyses

Gene and KEGG (Kyoto Encyclopedia of Genes and Genomes) pathway ontology enrichments were calculated using the R/Bioconductor package *missMethyl* (version 1.8.0, *gometh* function)[78], which performs one-sided hypergeometric tests taking into account and correcting for any bias derived from differing numbers of probes per gene interrogated by the array (see Additional file 13: Table S12). The annotation databases that were interrogated are http://www.kegg.jp/kegg/rest/keggapi.html for KEGG ontology, and the R/Bioconductor package *GO.db* (version 3.4.1) for gene ontology purposes. Appropriate backgrounds of total probes for each given context were employed in the corresponding analyses.

### Differential gene expression analyses

Gene expression data corresponding to RNAseq HTSeq-Counts from the GDC TCGA Kidney Clear Cell Carcinoma (KIRC) cohort were obtained from UCSC Xena Public Data Hub (http://xena.ucsc.edu/, dataset ID: TCGA-KIRC/Xena_Matrices/TCGA-KIRC.htseq_counts.tsv). Samples were filtered in order to fulfil the criteria of using only those cases with paired DNA methylation and gene expression data. Log2(count+1) data was further transformed to obtain integer count reads per gene condition. In order to strengthen downstream analyses, we removed non-variable and low-expressed genes (sum of expression across all the samples less than 1000 counts) to reduce the number of non-informative conditions. Differential expression analyses in aging (young versus old, groups 1 and 2 versus 5 respectively) or in cancer (Primary Tumor versus Solid Tissue Normal) conditions were performed with the R/Bioconductor package DESeq2 (version 1.16.1)[79]. DESeq2 analysis was performed with the default workflow and parameters. Genes were considered as differentially expressed if they satisfied *P* <0.05 after adjustment for multiple testing.

### Correlation analyses

RNAseqV2 log2(RSEM+1) normalized level 3 TCGA gene expression data were obtained for kidney normal tissue (KIRC) via UCSC Xena Public Data Hub (http://xena.ucsc.edu/). Samples were filtered so as to use only those with paired DNA methylation data. Non-variable and low-expression genes (those with sum of expression between all the samples of less than 10) were discarded. After filtering, pairwise Spearman correlations between DNA methylation level and gene expression level were calculated for all the combinations of probes and genes in normal kidney tissue samples, using probes that were previously detected as differentially hyper- and hypomethylated in cancer and aging.

## Ethics approval and consent to participate

The human data used in this study adhere to the ethical policies established by the TCGA consortium (Ethics, Law and Policy Group), the National Cancer Institute (NCI) and the National Human Genome Research Institute (NHGRI).

## Availability of data and materials

All data generated during this study are included in this published article and its supplementary information files, and are also available in the Zenodo public repository, DOI: 10.5281/zenodo.1086491, https://doi.org/10.5281/zenodo.1086491.

## Authors’ contributions

M.F.F., A.F.F. and G.F.B. conceived, coordinated and supervised the study. M.F.F, R.F. and J.R.T. designed all aspects of the research work. R.F. and J.R.T. collected the data and performed computational analyses. M.F.F, A.F.F, R.F. and J.R.T. wrote the manuscript. All authors revised, read, and approved the final manuscript.

## Acknowledgements

We are grateful to the members from the Cancer Epigenetics lab (FINBA, IUOPA) for their positive feedback and to Ronnie Lendrum for manuscript editing.

